# Whole genome sequencing of *Neisseria meningitidis* W isolates from the Czech Republic recovered in 1984 – 2017

**DOI:** 10.1101/346072

**Authors:** Michal Honskus, Zuzana Okonji, Martin Musilek, Jana Kozakova, Pavla Krizova

## Abstract

**Introduction:** The study presents the analysis of whole genome sequence (WGS) data for *Neisseria meningitidis* serogroup W isolates recovered in the Czech Republic in 1984 – 2017 and their comparison with WGS data from other countries.

**Material and Methods:** Thirty-one Czech *N. meningitidis* W isolates, 22 from invasive meningococcal disease (IMD) and nine from healthy carriers were analysed. The 33-year study period was divided into three periods: 1984-1999, 2000-2009, and 2010-2017.

**Results:** Most study isolates from IMD and healthy carriers were assigned to clonal complex cc22 (n = 10) in all study periods. The second leading clonal complex was cc865 (n = 8) presented by IMD (n = 7) and carriage (n = 1) isolates that emerged in the last study period, 2010 – 2017. The third clonal complex was cc11 (n = 4) including IMD isolates from the first (1984 – 1999) and third (2010 – 2017) study periods. The following clonal complex was cc174 (n = 3) presented by IMD isolates from the first two study periods, i.e. 1984 – 1999 and 2000 – 2009. One isolate of each cc41/44 and cc1136 originated from healthy carriers from the second study period, 2000 - 2009. The comparison of WGS data for *N. meningitidis* W isolates recovered in the Czech Republic in the study period 1984 – 2017 and for isolates from other countries recovered in the same period showed that clonal complex cc865, ST-3342 is unique to the Czech Republic since 2010. Moreover, the comparison shows that cc11 in the Czech Republic does not comprise novel hypervirulent lineages reported from both European and non-European countries. WGS data for Czech serogroup W meningococci point to the presence of MenB vaccine antigen genes and confirm the hypothesis about the MenB vaccine potential against *N. meningitidis* serogroup W. All 31 study isolates were assigned to Bexsero^®^ Antigen Sequence Types (BAST), and seven of them were of newly described BASTs.

**Conclusions:** WGS analysis contributed considerably to a more detailed molecular characterization of *N. meningitidis* W isolates recovered in the Czech Republic over a 33-year period and allowed for a spatial and temporal comparison of these characteristics between isolates from the Czech Republic and other countries. In addition, the WGS data precised the base for the update of the recommendation for vaccination in the Czech Republic.

## Introduction

The first global epidemic of invasive meningococcal disease (IMD) caused by the bacterium *Neisseria meningitidis* of serogroup W occurred in 2000 after the Hajj pilgrimage to Mecca, with cases reported in pilgrims and their close contacts from a number of countries [1]. This outbreak was due to the hypervirulent clonal complex cc11 of *N. meningitidis* W, designated the Hajj lineage [2]. After the Hajj epidemic, strains of the same clonal complex caused further outbreaks in African and South American countries [3].

A number of countries have recently reported serogroup W IMD cases caused by the hypervirulent clonal complex cc11 to be on the rise. The study of these isolates by the whole genome sequencing (WGS) method revealed two genetically close lineages: one is linked to the Hajj 2000 epidemic and its subsequent spread throughout the world, including to South Africa, and the other is recently reported from Latin America, England, and other countries. The European isolates serogroup W cc11 of the latter lineage are classified into two sub-lineages: original UK strain and novel 2013 UK strain [4, 5, 6, 7, 8, 9, 10]. In 2015-2016, the resurgence of *N. meningitidis* W cc11 was reported in Madagascar. Molecular characterization of isolates suggests local transmission of a single genotype [11]. Outbreaks of IMD caused by serogroup W cc11 were also reported in Australia in 2013-2015. The WGS analysis identified the original UK strain as the cause of these outbreaks [12, 13].

The incidence of meningococcal meningitis has been reported in the Czech Republic since 1943. IMD (including meningococcal meningitis) has been monitored within the national surveillance programme since 1993. The national case definition of IMD is in line with the European case definition from 2012. Isolates from 60 – 80 % of reported IMD cases are referred to the National Reference Laboratory for Meningococcal Infections in Prague (NRL) from all over the Czech Republic for confirmation and molecular characterization. In recent years, the proportion of IMD cases with the pathogen confirmed by the non-culture PCR method is on the rise (20 – 30 %). Serogroup B was prevailing most of the time while C was the leading serogroup in some years only. *N. meningitidis* of serogroup W is the cause of a low proportion of IMD cases in the Czech Republic but is associated with a high case fatality rate. It is important to monitor molecular characteristics of serogroup W isolates given the reported rise in IMD caused by hypervirulent complex cc11 of serogroup W in several countries and its ability to spread rapidly.

This study presents the first results of the WGS analysis of *N. meningitidis* W isolates from the Czech Republic recovered in 1984 – 2017.

## Material and methods

### Bacterial isolates and DNA extraction

All isolates of *N. meningitidis* W available in the NRL collection were analysed by whole genome sequencing. The NRL collection comprises about 5500 *N. meningitidis* isolates from IMD and healthy carriers deposited since 1971, along with their detailed characteristics and respective epidemiological and clinical data. Serogroup W isolates only represent a small proportion of strains in the NRL collection (1.24 %). The first available IMD isolate of *N. meningitidis* W is from 1984. The study period 1984 – 2017 was divided into three intervals to reflect the gradual increase in the proportion of serogroup W isolates among the total of IMD isolates: 1984 – 1999 (0.55 %), 2000 – 2009 (1.09 %), and 2010 – 2017 (4.31 %). A total of 31 isolates were selected for WGS analysis: five IMD isolates from 1984 – 1999, 13 isolates (six from IMD and seven from healthy carriers) from 2000 – 2009, and 13 isolates (11 from IMD and two from healthy carriers) from 2010 – 2017.

The bacterial cultures stored at -80 °C (Cryobank B, ITEST) were plated on chocolate Mueller-Hinton agar and cultured at 37° C and 5% CO_2_ for 18 – 24 hours. The isolates were assigned to serogroups by conventional serological methods (Pastorex Meningitidis Bio-RAD, antisera *N. meningitidis* ITEST, Bio-RAD) and confirmed by RT-PCR. The following step was the isolation of deoxyribonucleic acid (DNA) using the QIAamp DNA Mini Kit (QIAGEN) according to the manufacturer’s instructions.

### Whole genome sequencing and WGS data processing

The whole genome sequencing of isolates of *N. meningitidis* W was conducted by the European Molecular Biology Laboratory (EMBL), Heidelberg, Germany. The Illumina MiSeq platform was used for sequencing against the reference genome sequence of *N. meningitidis* strain MC58. The result was overlapping sequences approximately 300 bp in length. WGS data were subsequently processed using the Velvet *de novo* Assembler software. To optimise the procedure, the Velvet-Optimiser script was used [14]. The K-mer length parameter varied between isolates from 91 to 183 (151 on average). The resultant genome contigs were submitted to the Neisseria PubMLST database (www.pubmlst.org/neisseria/), which runs the BIGSdb (Bacterial Isolate Genome Sequence Database) platform [15, 16], under the following IDs: 38989, 41191, 57208, 57209, 57211 – 57227, 57829, 57832, 57834, 57836, 57841 – 57846.

### Genome analysis and WGS data visualization

In the PubMLST database, the genome contigs of individual isolates were automatically scanned and characterized by allelic profile of the genes, which are determined in the NRL by conventional sequencing methods (*abcZ*, *adk*, *aroE*, *fumC*, *gdh*, *pdhC*, *pgm*, *porA*, *fetA*, *nhba*, *nadA*, and *fhbp*). Based on the allelic profile of seven MLST genes, isolates were assigned to sequence type (ST) and clonal complex (cc) [17]. Allelic variants were determined in variable regions (VR) contained in the *porA* (twice) and *fetA* (once) genes. Each unique combination of such allelic variants is called a finetype [18]. Furthermore, allelic and peptide variants of MenB vaccine antigens (*nhba*, *nadA*, and *fhbp*) were determined [19, 20, 21, 22, 23]. A Bexsero^®^ antigen sequence type (BAST) is a unique combination of peptide variants of these genes and allelic variants of two *porA* gene variable regions [24]. New gene and peptide variants were scanned manually, added to the database, annotated, and numbered using the automated data entry tool of the BIGSdb platform.

Genomes were analysed and compared using the BIGSdb Genome Comparator tool, which is part of the PubMLST database [16]. WGS data for isolates were compared using the core genome cgMLST scheme v1.0 for *N. meningitidis* - (1605 loci) [25].

The distance matrices, which are based on the number and allelic variability of the genes contained in individual schemes, were generated automatically and phylogenetic networks were constructed using the SplitsTree4 software which uses the NeighborNet algorithm [26]. Phylogenetic analysis results were edited graphically by the Inkscape tool (www.inkscape.org/en/). Isolates are coloured according to detection year (yellow 1984 - 1999, green 2000 – 2009, and red for 2010 – 2017).

### Comparison of WGS data between isolates of *N. meningitidis* W from the Czech Republic and other countries

To gain a more detailed insight into the genetic diversity of the Czech isolates of serogroup W *N. meningitidis*, we compared WGS data between countries, which facilitates the study of the genetic profile of the population of Czech isolates and of the relationship between their genetic diversity and geographical distribution. Using the data from the PubMLST database, a selection was made of all available European and non-European *N. meningitidis* serogroup W isolates belonging to two clonal complexes (cc11 and cc22) that are the most widespread worldwide. Only isolates for which full MLST profiles and WGS data were available (sequence bin size >= 2 Mbp) were included in the study. These criteria were met by 1094 cc11 and 159 cc22 isolates from other countries.

The overall genetic diversity within serogroup W isolates is shown in three phylogenetic networks. Each of them represents the comparison between Czech serogroup W isolates and those from other countries, which were available in the PubMLST database. Group 1 consists exclusively of UK isolates (n = 901). Group 2 comprises isolates from continental European countries (n = 399): France (n = 136), the Netherlands (n = 110), Sweden (n = 67), Italy (n = 41), Ireland (n = 21), Finland (n = 10), Portugal (n = 4), Germany, Island, and Malta (two isolates from each), Greece, Croatia, Norway, and Spain (one isolate from each). Group 3 includes non-European isolates (n = 363): from the Republic of South Africa (n = 130), Canada (n = 75), Niger (n = 37), China (n = 25), Burkina Faso (n = 17), Cameroon (n = 12), Madagascar (n = 8), Turkey (n = 7), Algeria (n = 7), Mali (n = 6), USA (n = 5), Russia (n = 4), and Senegal (n = 4), two isolates from each Japan, Morocco, Saudi Arabia, Togo, Benin, and Chad, and one isolate from each Mauritius, Djibouti, and Central African Republic. Eleven isolates with missing data on the place of detection were added to this group. For all of these isolates, WGS data (sequence bin size >= 2 Mbp) and full MLST profiles were available.

Isolates from the Czech Republic and other countries were compared using the Genome Comparator tool at the cgMLST level (1605 loci). In the phylogenetic networks, isolates are coloured according to detection year. Isolates recovered before 2000 are highlighted in yellow, isolates from 2000 – 2009 in green, and isolates from 2010 – 2017 in red. The isolates from other countries with missing detection year are highlighted in grey. The study isolates from the Czech Republic are marked with squares coloured according to the study intervals and numbers under which they are registered in the NRL collection of *N. meningitidis* isolates.

## Results

### Distribution of *N. meningitidis* W isolates from the Czech Republic and assignment to clonal complexes

First figure (Fig. 1) shows the distribution of all (part A) or IMD (part B) *N. meningitidis* serogroup W isolates from the Czech Republic by study interval. Clonal complex affiliation of isolates is highlighted in colours. Five isolates from 1984 – 1999 are exclusively from IMD and belong to three clonal complexes (cc11, cc22, and cc174), with cc11 being predominant (60 %). In 2000 – 2009, more cc22 isolates were recovered (two IMD isolates and three carriage isolates). In that period, complex 174 is represented by two IMD isolates, and three isolates were unassigned to clonal complex (ccUA) (two IMD isolates and one carriage isolate). In 2000 – 2009, three carriage isolates belonging to three different clonal complexes, cc41/44, cc53, and cc1136, were recovered. No cc11 isolate was registered in the Czech Republic in that period. In 2010 – 2017, cc22 isolates can be seen again (three IMD isolates and one carriage isolate); one IMD cc11 isolate and eight cc865 isolates emerged (seven IMD isolates and one carriage isolate).

**Figure 1:**
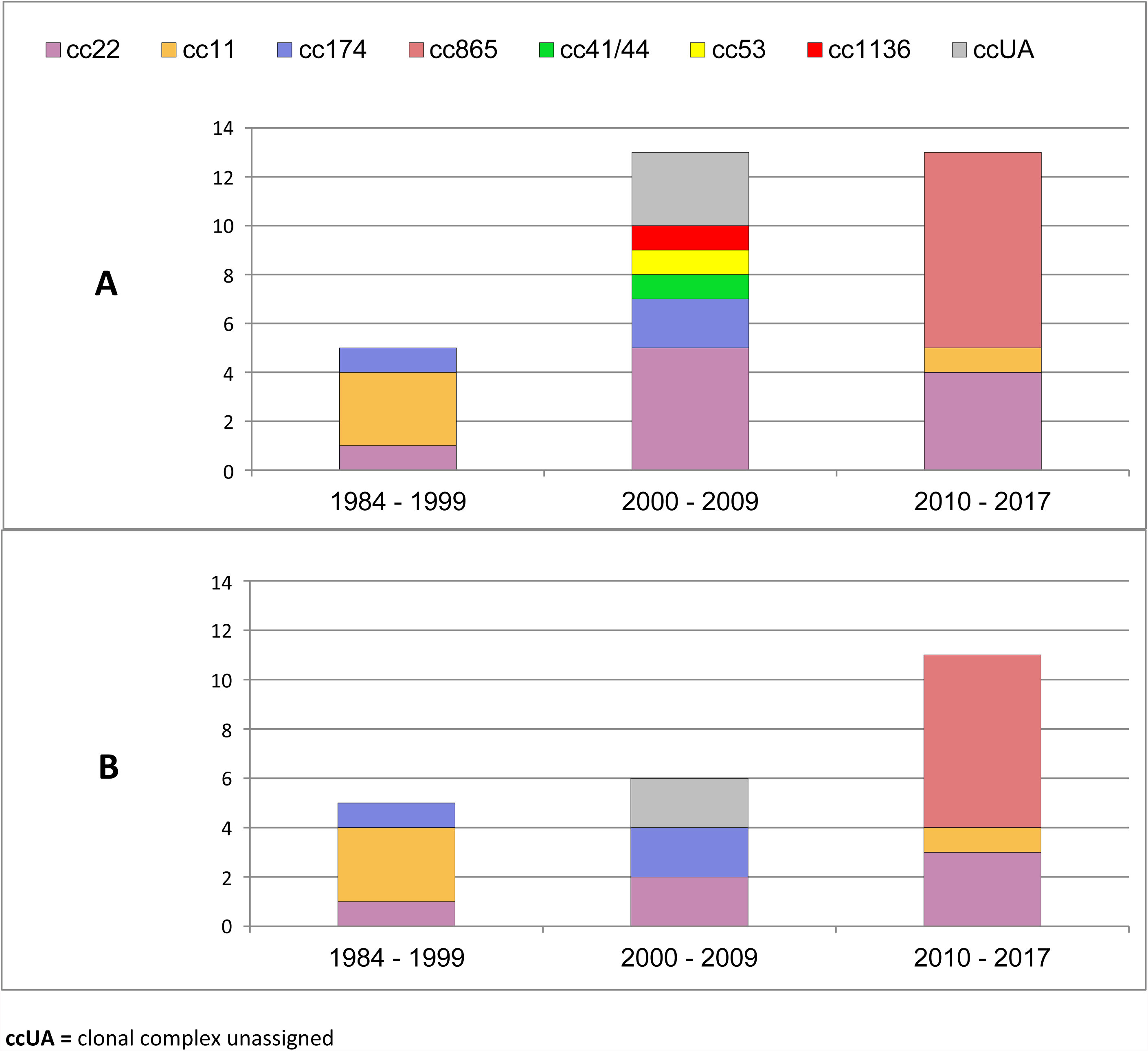
Distribution of *N. meningitidis* serogroup W isolates from Czech Republic in three time periods and assignment to clonal complexes. Isolates were recovered from 1984 to 2017. Part A: all isolates (n = 31), part B: only isolates from invasive meningococcal disease (n = 22).

Conclusion: IMD cc22 isolates were recorded throughout the all study periods, while cc11 was only found in 1984 – 1999 and 2010 – 2017, and cc174 in 1984 – 1999 and 2000 - 2009. The most relevant finding is a high incidence of cc865 isolates in the last study period.

### Genetic relationships between *N. meningitidis* W isolates from the Czech Republic

The generated phylogenetic network confirmed that most (81 %; n = 25) *N. meningitidis* W isolates from the Czech Republic belong to four clonal complexes: cc22 (n = 10), cc865 (n = 8), cc11 (n = 4), and cc174 (n = 3). All these clonal complexes are clearly delineated in the phylogenetic network (Fig. 2).

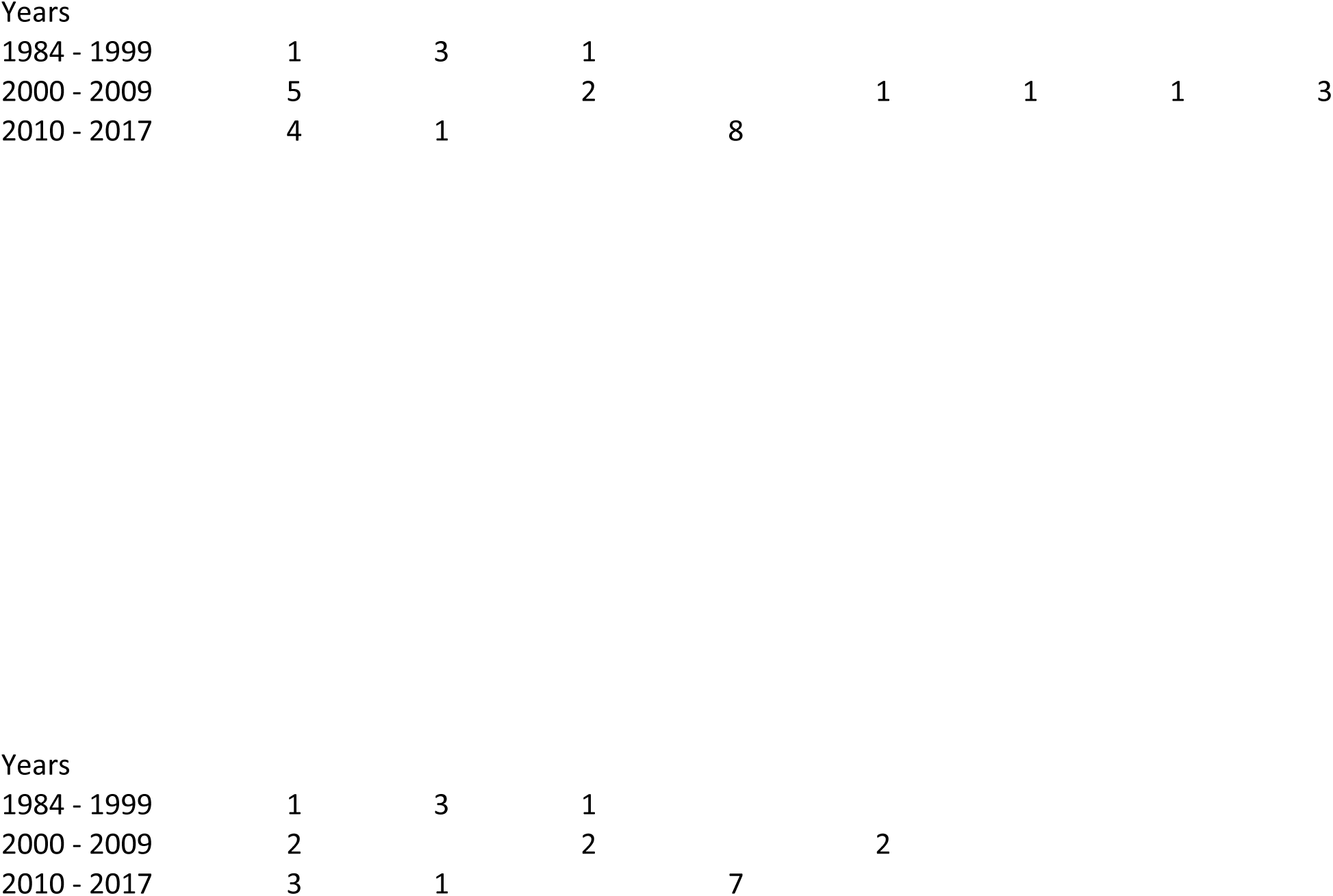

The clonal complex cc865 shows the highest homogeneity, which is consistent with the fact that all isolates of this clonal complex belong to the same sequence type ST-3342 and were recovered in the most recent interval, i.e. 2010 – 2017 (one in 2011, two in 2012, three in 2016, and two in 2017). Based on the data available in the PubMLST database, cc865 is uncommon in serogroup W and was only detected in seven countries (one isolate from each Germany, Spain, the Netherlands, Greece, Romania, Russia, and the Republic of South Africa). So far, sequence type ST-3342 has only been identified in the Czech Republic. All cc865 isolates from other countries (n = 7) were assigned to different sequence types. It is interesting to note that each of these cc865 isolates has a unique sequence type (ST-1232, ST-6444, ST-8172, ST-8608, ST-10799, ST-11589, and ST-12256). Therefore, it can be assumed that cc865 ST-3342 isolates from the Czech Republic originate from a common ancestor that has recently evolved in the country. The ongoing diversification of the *N. meningitidis* W cc865 ST-3342 population in the Czech Republic is demonstrated by gene changes in MenB vaccine antigens (*fhbp*, *nhba*, and *nadA*). Although all ST-3342 isolates contain a peptide variant of the *nhba* 89, two isolates from 2017 carry a same synonymous point mutation that switches allelic variant 257 to variant 1438 (Tab. 1).

**Table 1:**
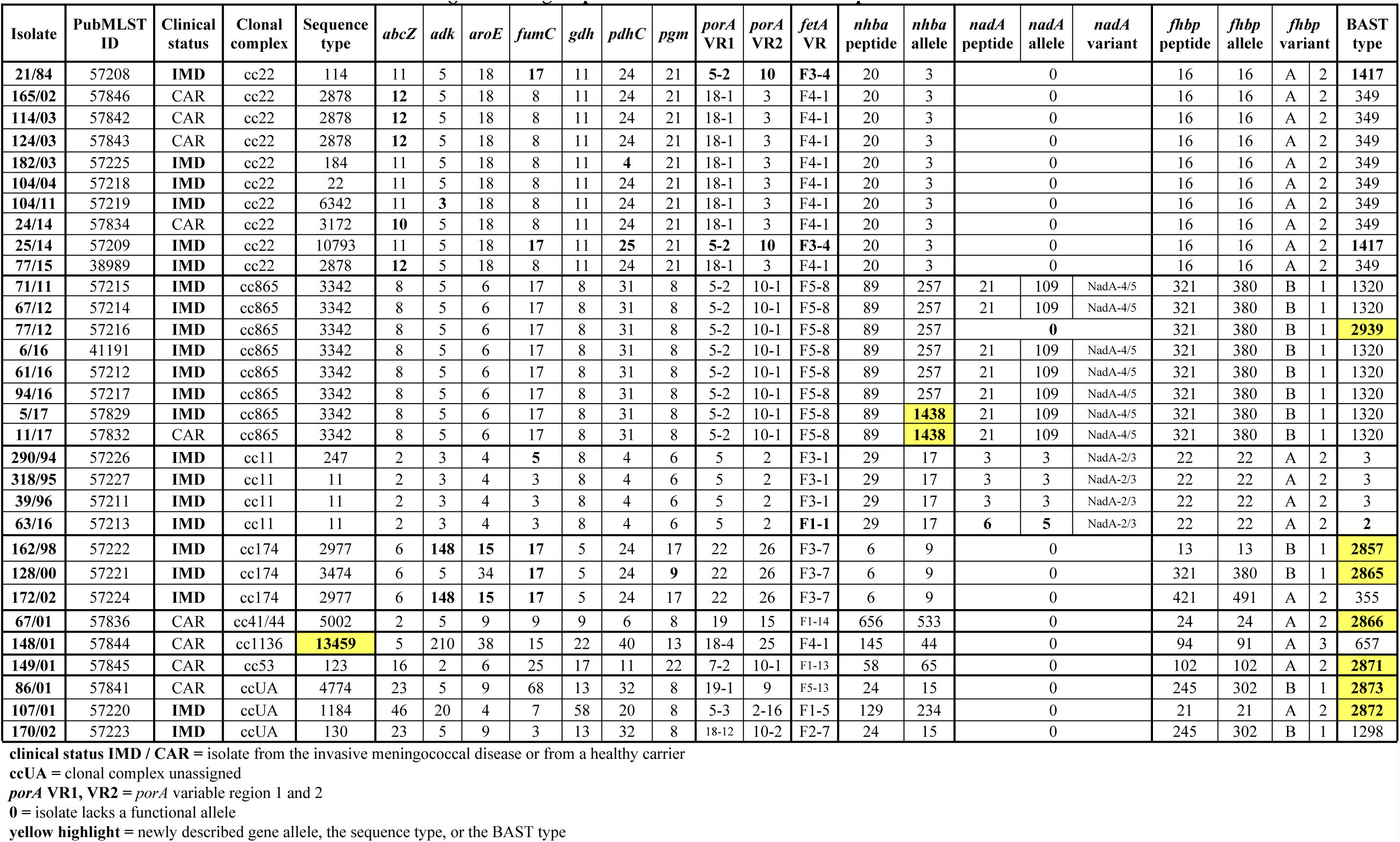
Molecular characterization of 31 *N. meningitidis* serogroup W isolates from the Czech Republic recovered from 1984 to 2017.

This allelic form of the *nhba* gene has not yet been known, and its sequence was submitted to the PubMLST database to be assigned a new allele number. Isolate 77/12 has lost a functional allele of the *nadA* gene which is present in all other isolates (allele 109, peptide variant 21), as also reflected by assignment to a different BAST type – 2939, with all other isolates being classified into BAST 1320. The *fhbp* gene in all isolates is represented by allele 380 (peptide ID 321, subfamily 1/B). All cc865 ST-3342 isolates from the Czech Republic have the same finetyping antigens (5-2,10-1:F5-8).

Ten out of the 31 study Czech isolates of *N. meningitidis* W belonged to clonal complex cc22. The higher diversity of the phylogenetic network is reflected in the fact that these 10 isolates are assigned to seven different sequence types: ST-2878 (n = 4), ST-22, ST-114, ST-184, ST-3172, ST-6342, and ST-10793 (Tab. 1). In the Czech Republic, cc22 isolates were recovered in all three study intervals. Isolates 21/84 (ST-114) and 25/14 (ST-10793) are separated from other isolates and share similar molecular characteristics despite the large time gap between their detection (1984 vs. 2014). Unlike all other cc22 isolates (18-1,3:F4-1; BAST 349), they have the same difference in finetyping antigens and BAST type (5-2,10:F3-4; BAST 1417), and unlike all other isolates with allele 8, they exhibit the same allele change of one MLST gene, *fumC*, to allele 17. Isolate 104/04 (ST-22) is also partly separated from all other cc22 isolates. Three isolates of ST-2878 (165/02, 114/03, and 124/03) show high relatedness. The fourth isolate, 77/15, of the same ST (ST-2878) is evolutionarily more distant, probably as a result of the accumulation of genetic changes due to the large time gap between their detection. All cc22 isolates show full homogeneity in the MenB vaccine antigen genes. Characteristics of the MenB vaccine antigen genes: *nhba* – allele 3 (peptide variant 20), *fhbp* – allele 16 (peptide ID 16, subfamily 2/A), and absence of a functional form of *nadA*.

Only four study Czech isolates belonged to the hypervirulent clonal complex cc11, which is not consistent with the recent global upward trend in *N. meningitidis* W cc11 cases. Three of these four cc11 isolates were recovered between 1994 and 1996 and thus do not belong to the new lineages of *N. meningitidis* W cc11, which are spreading worldwide. Except the fact that isolate 290/94 was assigned to ST-247 (the other two are of ST-11; the difference in the allele of the *fumC* gene – ST-11 allele 3 vs. ST-247 allele 5), these isolates share identical molecular characteristics: finetyping antigens 5,2:F3-1; *nhba* – allele 17 (peptide variant 29), *fhbp* – allele 22 (peptide ID 22, subfamily 2/A), *nadA* – allele 3 (peptide variant 3), and BAST type – 3 (Tab. 1). The molecular characteristics of isolate 63/16 from 2016 are consistent with those of the new lineages of *N. meningitidis* W cc11, but this IMD isolate originates from a Canadian traveller from Hungary to the Czech Republic. Unlike the three previous isolates, isolate 63/16 shows changes in finetyping antigens (5,2:F1-1) and the *nadA* gene (allele 5, peptide variant 6), as also reflected in BAST type changed to BAST 2.

Three isolates belong to clonal complex cc174. Isolates 162/98 and 172/02 are assigned to sequence type ST-2977 and isolate 128/00 to ST-3474, as is reflected by their positions in the phylogenetic network. These sequence types differ in three alleles of the genes *adk*, *aroE*, and *pgm* (MLST). All these isolates recovered between 1998 and 2002 share identical molecular characteristics in terms of finetyping antigens (22,26:F3-7), the *nhba* gene (allele 9, peptide variant 6), and absence of a functional allele of the *nadA* gene. Each isolate has a unique allele of the *fhbp* gene and unique BAST (Tab. 1).

Three clonal complexes, each represented by one isolate, i.e. 67/01 (cc41/44, ST-5002), 149/01 (cc53, ST-123), and 148/01 (cc1136, ST-13459), are clearly delineated from other clonal complexes and evolutionarily distant from each other. Isolate 148/01 showed as yet undescribed MLST gene allele combination. Using the PubMLST database, this combination was assigned a new sequence type ST-13459.

Three isolates unassigned to clonal complex by the PubMLST database are 170/02 (ST-130), 107/01 (ST-1184), and 86/01 (ST-4774). While isolate 107/01 is clearly separated in the phylogenetic network, isolates 86/01 and 170/02 show a relatively high level of relatedness. Although having different finetyping antigens (ST-130 finetyping 18-12,10-2:F2-7 vs. ST-4774 finetyping 19-1,9:F5-13), these two isolates only vary in a single MLST gene allele (gene *fumC* – ST-130 allele 3 vs. ST-4774 allele 68). Nevertheless, they are fully congruent in MenB vaccine antigen genes: *nhba* – allele 15 (peptide variant 24), *fhbp* – allele 302 (peptide ID 245, subfamily 1/B), and absence of a functional form of *nadA* (Tab. 1).

### Genetic diversity of *N. meningitidis* W cc11 isolates

The complex phylogenetic network of worldwide cc11 isolates (Fig. 3) clearly shows that Czech isolates 290/94, 318/95, and 39/96 do not belong to the new *N. meningitidis* W cc11 lineages that now cause IMD worldwide. These three isolates belong to genetically distant lineages grouping mostly isolates recovered before the year 2000. It was also confirmed that the imported isolate 63/16, on the contrary, belongs to these modern hypervirulent lineages. It forms a large cluster along with many isolates recovered almost exclusively after 2010. This cluster is clearly divided into several subpopulations, which is in line with studies from other European countries. The information presented thus confirms that the recent increase in IMD caused by new hypervirulent serogroup W cc11 lineages, as observed in other European and non-European countries, still did not reach the Czech Republic.

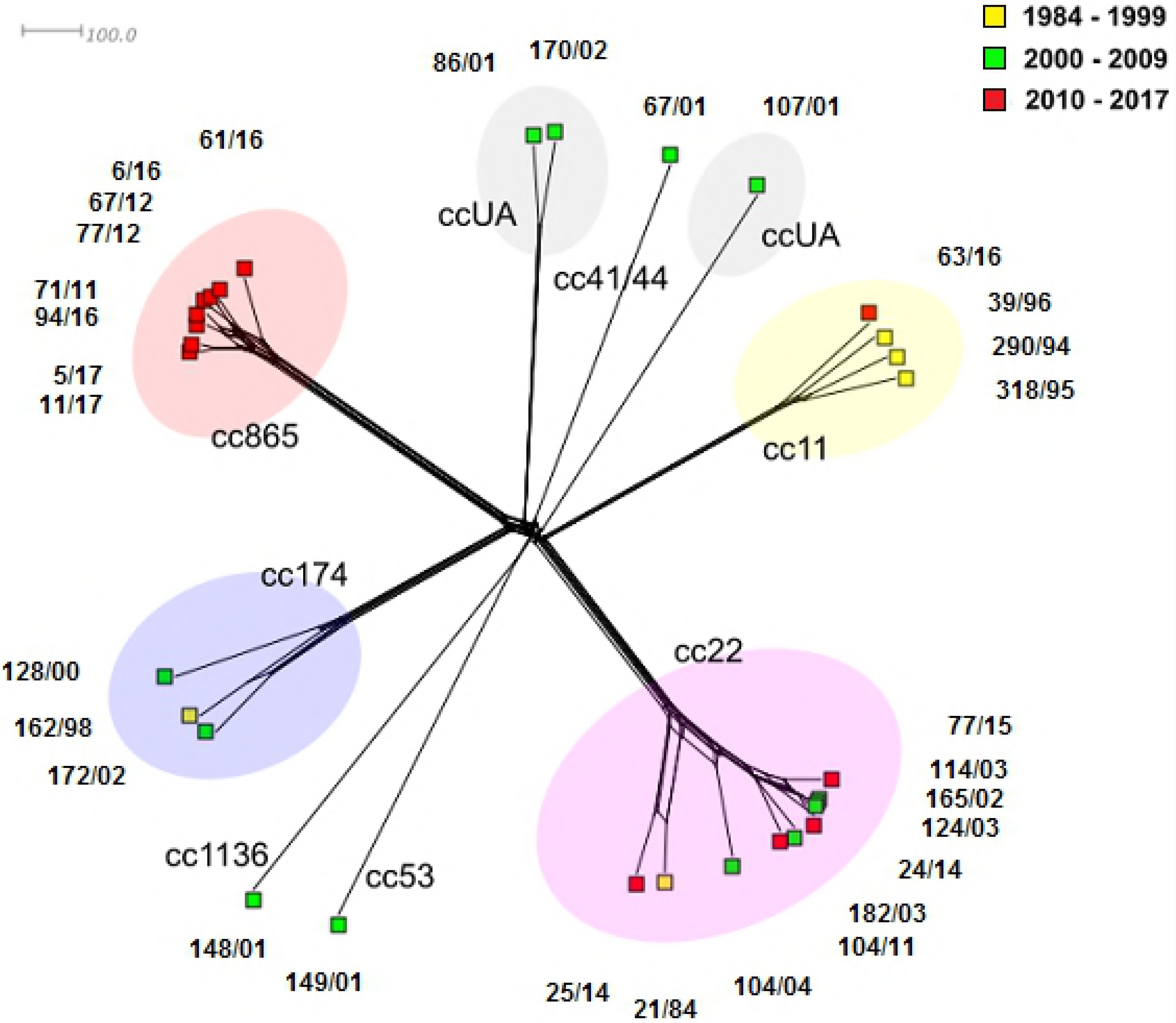

### Genetic diversity of *N. meningitidis* W cc22 isolates

A relatively even distribution of Czech isolates can be seen in the phylogenetic network of clonal complex cc22 (Fig. 4), which is consistent with the high variability of sequence types within this complex. An exception is a cluster of four isolates of ST-2878 (165/02, 114/03, 124/03, and 77/15), as is expected given their assignment to the same sequence type. Isolate 24/14 (ST-3172) also appears to be closely related to ST-2878 isolates. As can be seen from the molecular characteristics (Tab. 1), the only difference between these sequence types is in the allele of the *abcZ* gene (ST-2878 – allele 12 vs. ST-3172 – allele 10). This was also confirmed by the described genetic relatedness of isolates 21/84 and 25/14. It appears that in European countries other than the Czech Republic, cc22 is less common than cc11. The proportion of cc22 among isolates from the Czech Republic (32 %) is uncommonly high in comparison with other European countries.

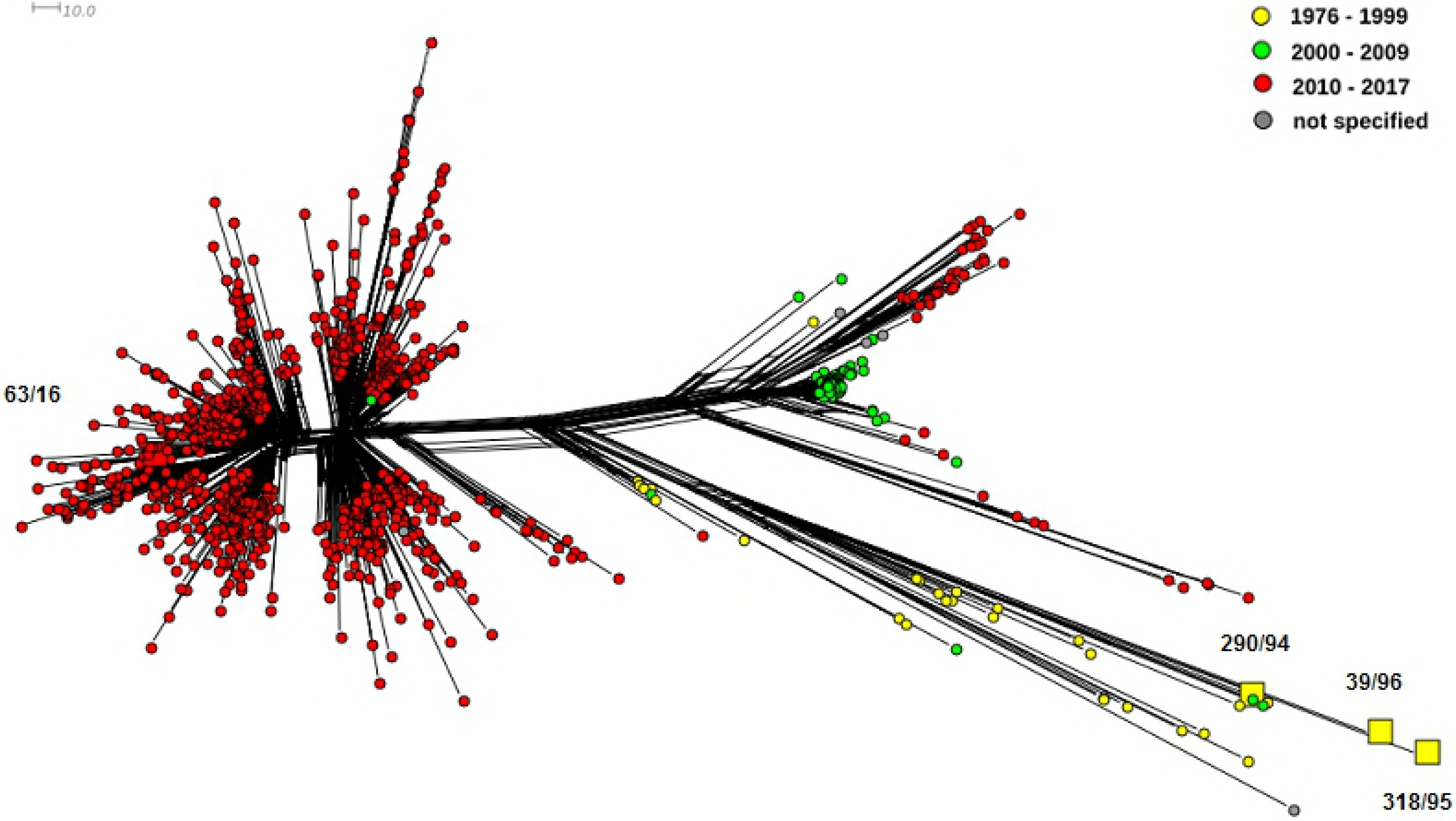

### Genetic relationships between *N. meningitidis* W isolates from Czech Republic and United Kingdom

The phylogenetic network, which represents the genetic diversity of isolates from the Czech Republic and United Kingdom (Fig. 5), shows that most serogroup W isolates belong to two clonal complexes, cc11 and cc22. The cc11 isolates are clearly more numerous than the cc22 isolates. In the phylogenetic network, these two groups are clearly distinct from each other and genetically distant from each other. The phylogenetic network of clonal complex cc11 displays several subpopulations of new W cc11 lineages, one genetically distinct subpopulation of isolates recovered mainly in 2000 – 2009, and several historical lineages from 1975 – 1999. As few as 30 (3 %) isolates from this selection were assigned to other clonal complexes. In this population of other clonal complexes where isolates from the Czech Republic account for more than half, it can be seen a rather heterogeneous cc174 lineage comprising both Czech and UK isolates and eight cc865 isolates originating exclusively from the Czech Republic. Clonal complex cc11 comprises one isolate, 63/16, from an imported case of IMD, which belongs among new W cc11 lineages, and three Czech isolates (290/94, 318/95 a 39/96) recovered before the year 2000 and belonging to clearly separated and genetically distinct historical lineages. In conclusion, it can be stated that serogroup W clonal complexes from the Czech Republic and the UK differ in the population structure. In the UK, the cc11 lineages are predominant while cc22 isolates are rather rare and isolates belonging to other clonal complexes are found only sporadically. On the other hand, the clonal complex most often detected in the Czech Republic is cc22 (32 %), and isolates from other clonal complexes (almost 55 %) are also common.

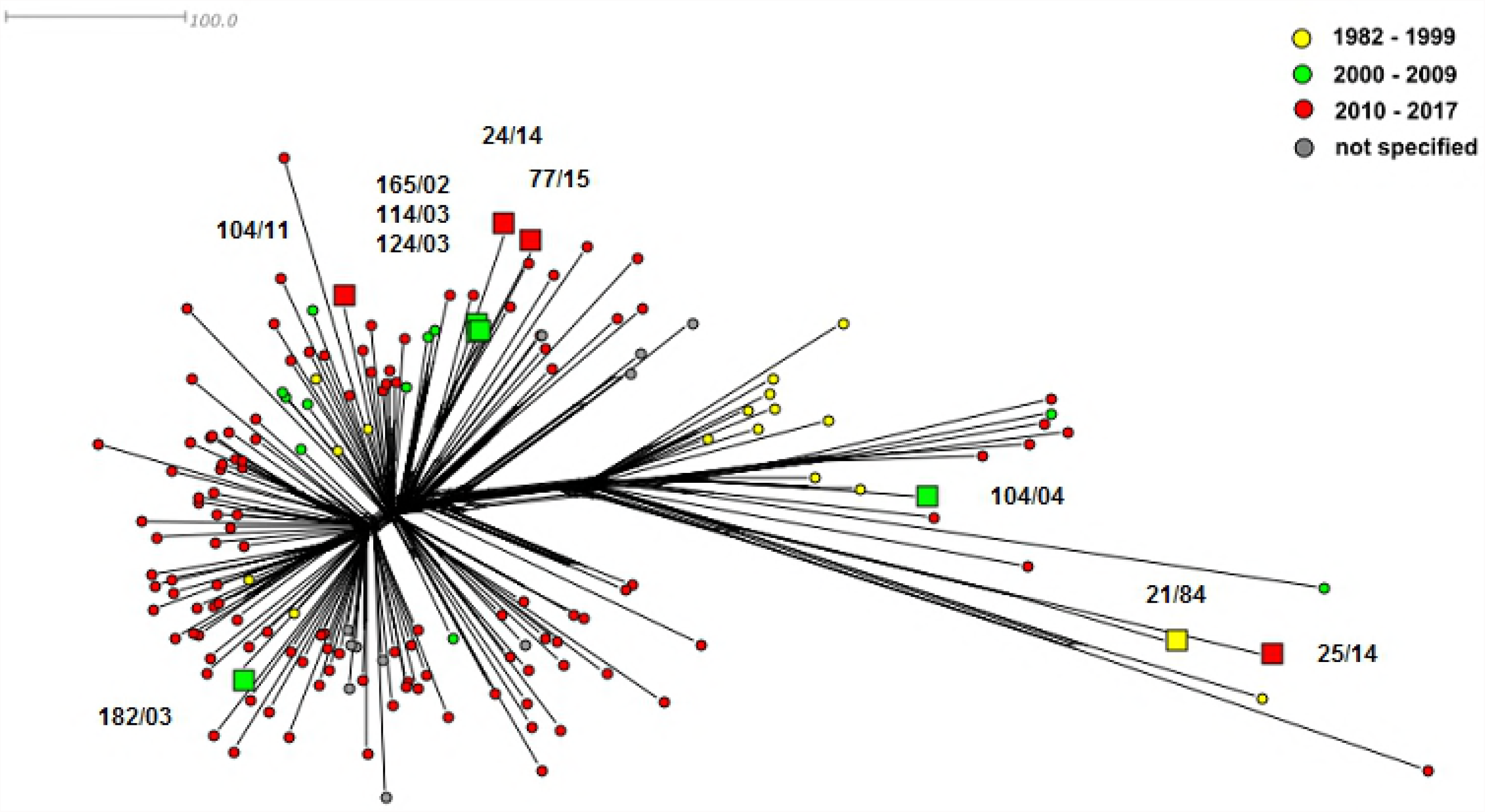

### Genetic relationships between *N. meningitidis* W isolates from Czech Republic and continental Europe

The following phylogenetic network (Fig. 6) illustrative of the genetic variability of serogroup W isolates from the Czech Republic and other European countries shows more heterogeneity than the previous figure. Again, most isolates belong to clonal complexes cc11 and cc22, but unlike cc22 isolates, cc11 isolates experienced a considerable decline. It is evident that cc22 is more commonly detected in European countries than in the UK. Nevertheless, cc11 isolates are still more frequent than cc22 isolates. The proportion of isolates assigned to other clonal complexes is also higher. In the phylogenetic network, there can be seen a lineage of four ccUA isolates from the Netherlands from 2011 – 2016 showing partial relatedness to cc22 and a distinct lineage of two cc8 isolates from France from 1978. In a large heterogeneous group of isolates belonging to clonal complexes other than cc11 and cc22, a separate cc174 lineage appears again, comprising isolates from the Czech Republic and other European countries. Clonal complex cc865, so far only represented by isolates from the Czech Republic, was extended by one isolate (cc865, ST-12256) from the Netherlands from 2017. It is interesting to note that six isolates of sequence type ST-9316 unassigned to clonal complex from France (n = 5) and Ireland (n = 1) from 2015 – 2016 show a higher relatedness to the Czech cluster of cc865 (ST-3342) isolates than the Dutch isolate belonging to the same clonal complex.

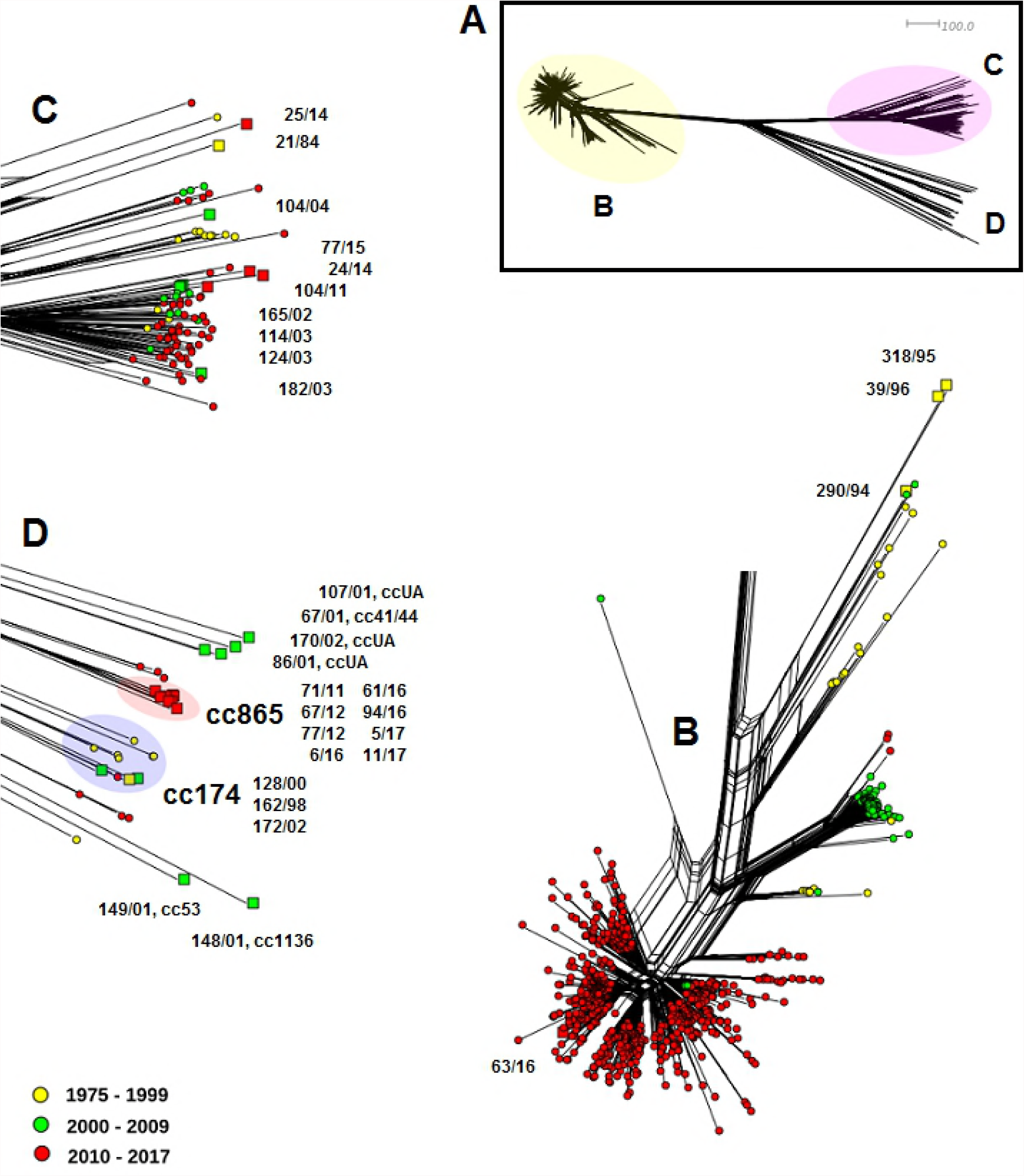

### Genetic relationships between *N. meningitidis* W isolates from Czech Republic and non-European countries

Similarly to the previous two figures (Fig. 5 and Fig. 6), two main clusters of cc11 and cc22 isolates can be seen in this figure (Fig 7). Both clusters, and cc11 in particular, show higher heterogeneity, which is probably due to the geographical diversity of the isolates. In clonal complex cc11, there are more isolates recovered before 2010 in comparison with two groups of isolates from European countries and the UK, where the two modern lineages were predominant (particularly among the European isolates). The proportion of cc22 isolates declined in comparison with the European isolates while the number of isolates belonging to other clonal complexes remained nearly unchanged, with many isolates originating from the Czech Republic again. This group comprises cc174 isolates from both the Czech Republic and non-European countries. An additional clonal complex cc175 can be seen, including nine highly related isolates from African countries (three from Niger, ST-2881; two from Benin, ST-2881; two from Togo, ST-2881, and two from Burkina Faso, ST-2881 and ST-8638) from 2003 – 2010 and one genetically more distant isolate (ST-6218) from the Republic of South Africa from 2003. Clonal complex cc865, comprising eight isolates from the Czech Republic (ST-3342), was added with one South African isolate (cc865, ST-8608) from 2009. Interesting to note are nine highly related cc4821 ST-8491 isolates from China. According to the data available from the PubMLST database, no serogroup W isolate belonging to cc4821 has not yet been reported by any other country, so, this clonal complex of serogroup W is endemic in China [27].

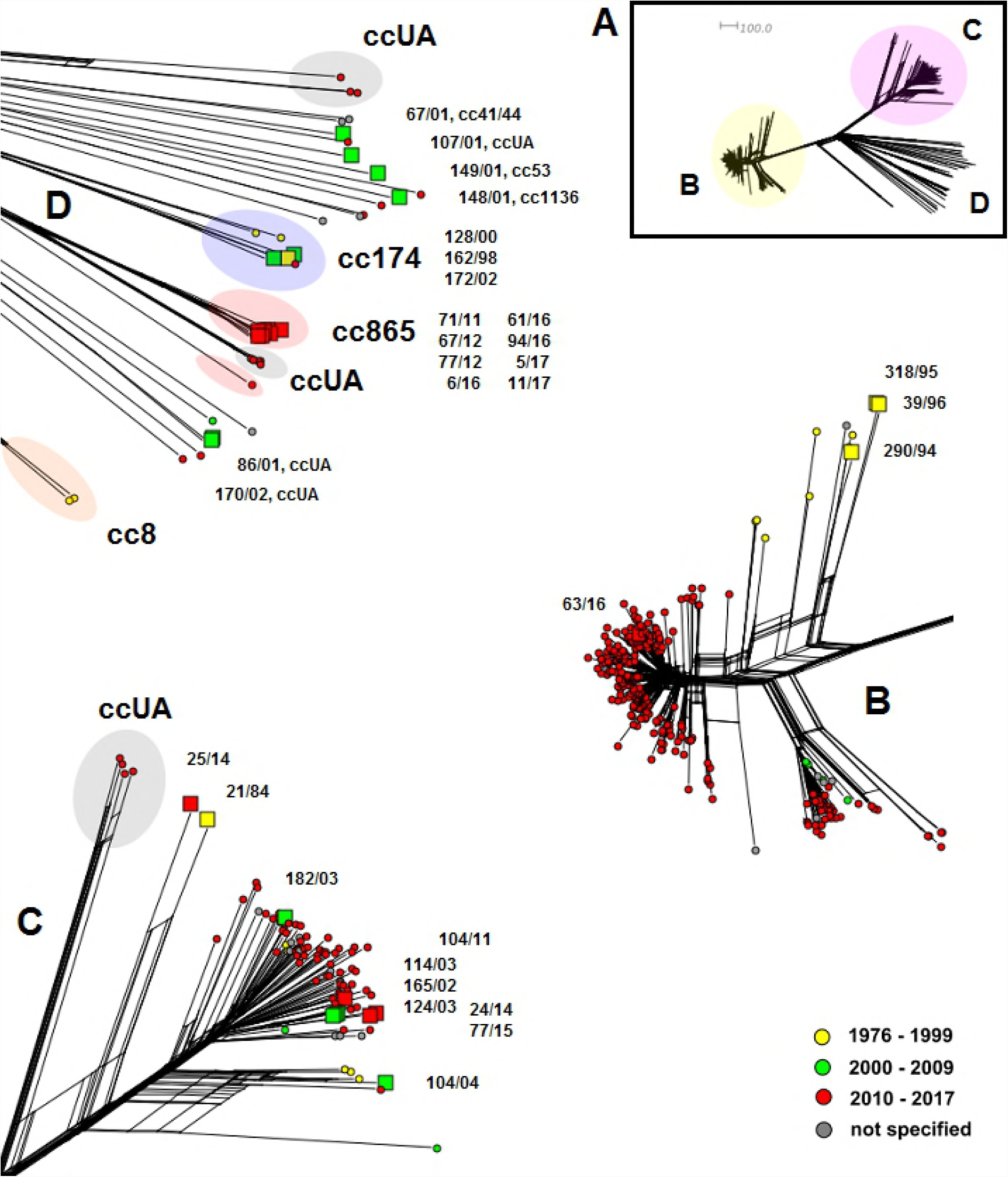

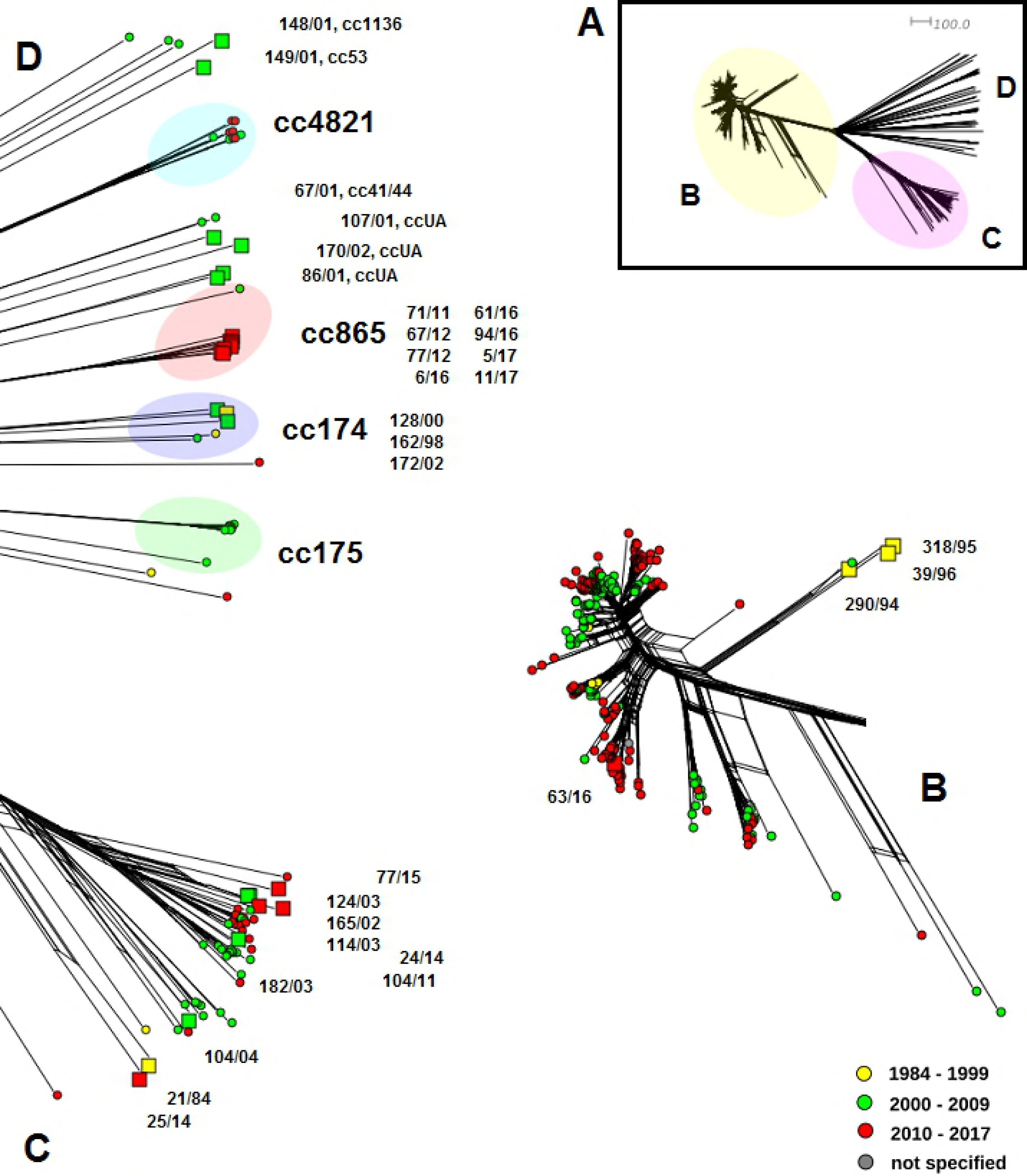

## Discussion

In countries of Sub-Saharan Africa, Middle East, and Western Europe or in Australia, cc11 is the serogroup W clonal complex which is on the rise. Recently, *N. meningitidis* W cc11 has even become the main cause of IMD in the UK, France, the Netherlands, and Sweden. At present, most cases of IMD in the UK, the Netherlands, Sweden, and France are caused by the lineages called the original UK strain and 2013-UK strain of hypervirulent *N. meningitidis* W cc11 [8, 9, 10, 28, 29]. Our WGS study shows that the Czech isolates of *N. meningitidis* W do not belong to these novel hypervirulent cc11 lineages. The potential for a rapid spread of hypervirulent *N. meningitidis* W cc11 in the world was demonstrated in connection with the World Scout Jamboree held in Japon in 2015, with cases of IMD caused by *N. meningitidis* W cc11 reported in jamboree participants and their close contacts (four cases in Scotland and two cases in Sweden) [30]. Given the increased international travel, surveillance of this hypervirulent complex in the Czech Republic is of high relevance. Using the WGS method, one IMD isolate of *N. meningitidis* W was confirmed to belong to the novel hypervirulent lineage cc11. This isolate originated from an imported case of IMD in a traveller of Canadian nationality who came to the Czech Republic from Hungary in 2016.

The most interesting finding of this study is that eight of 31 Czech isolates of *N. meningitidis* W belong to clonal complex cc865, which is uncommon for serogroup W as can be inferred from the data available in the PubMLST database. All Czech cc865 isolates are genetically highly homogeneous, were isolated exclusively between 2010 and 2017, and belong to the same sequence type, ST-3342, which, to date, has only been reported from the Czech Republic (PubMLST). This body of evidence supports the assumption that isolates cc865, ST-3342 originate from a common ancestor that evolved in the Czech Republic.

The WGS has a higher resolution in comparison with conventional sequencing methods and demonstrated the genetic heterogeneity of the population of *N. meningitidis* W cc11. The Hajj lineage continued to spread in the Middle East while in South African and meningitis belt countries, other strains of *N. meningitidis* W were recovered along with it. South America, the UK, and France share another genetically different strain of *N. meningitidis* W cc11 [31]. The WGS method demonstrated the diversification of *N. meningitidis* W in the African meningitis belt during the period 1994 – 2012 [32]. The study isolates belonged to cc11 (83 out of 92) or cc175 (nine out of 93). *N. meningitidis* W cc11 isolates were classified into four major subclades, I – IV, linked to specific epidemiological situations: subclades I and II were not linked to outbreaks, subclade II was linked to the 2002 outbreak in Burkina Faso, and subclade IV was linked to the 2000 outbreak in Saudi Arabia.

The WGS was introduced into molecular surveillance of IMD in the Czech Republic [33] in line with the ECDC strategy [34]. Our paper is the first presentation of the results of the WGS study of *N. meningitidis* W isolates from the collection of the Czech NRL spanning a 33-year period. The first clonal study of the historical collection of isolates of all serogroups recovered from cases of IMD in the Czech Republic over a more than 40-year period was based on the analysis of MLST results [35].

In response to the rise in IMD caused by hypervirulent lineages of *N. meningitidis* W cc11, immunisation campaigns using tetravalent meningococcal conjugate ACYW vaccine were launched in some countries, for example in Chile and the UK [3, 36]. In view of the low incidence of IMD and absence of hypervirulent lineages of *N. meningitidis* W cc11 in the Czech Republic, no vaccination campaign is considered in this country, and individual protection is recommended [37]. Further molecular surveillance of IMD is needed to assess the outcomes of the recommended vaccination strategy in the Czech Republic.

The UK study pointed out the potential of the MenB-4C vaccine against hypervirulent *N. meningitidis* W cc11 [38, 39]. Despite being licensed for the prevention of IMD caused by serogroup B, the MenB-4C vaccine contains antigens which are not serogroup B specific and can provide protection against other capsular serogroups, which share the same antigens. MenB vaccine is expected to elicit bactericidal antibodies against *N. meningitidis* W cc11 in children (infants and toddlers). The WGS detection of MenB vaccine genes in Czech *N. meningitidis* W isolates suggests the potential of this vaccine also against serogroup W isolates.

## Acknowledgment

Supported by the Ministry of Health of the Czech Republic, grant no. 15-34887A. All rights reserved. This publication made use of the PubMLST website (https://pubmlst.org/) developed by Keith Jolley (Jolley & Maiden 2010, BMC Bioinformatics, 11:595) and sited at the University of Oxford. The development of that website was funded by the Wellcome Trust.

